# Preparation and pharmacokinetics of genistein MePEG-PLGA copolymer micelles

**DOI:** 10.1101/620898

**Authors:** Mina Swartz, John Smith

**Affiliations:** Department of Biochemistry, University of South Carolina, Columbia SC, 29208

**Author notes:** Address: 631 Sumter Street, Columbia, SC 29208.

## Abstract

In this report, we demonstrated a novel technique to prepare genistein (GEN) MePEG-PLGA copolymer micelles. Initial stability and pharmacokinetic behavior in rats after intravenous administration were investigated. The micelles were prepared by modified self-emulsifying solvent evaporation method. The morphology, encapsulation efficiency, drug loading, particle size and Zeta potential were investigated. The release behavior was investigated by dynamic membrane dialysis technique. The micelles were stored in a refrigerator at 4 °C, and samples were taken after 1 d, 10 d, 1 month, 3 months, and 6 months, and the encapsulation efficiency and drug loading were examined. The GEN micelles were injected into the tail vein of healthy rats. The blood concentration of GEN in rats was determined by HPLC. The plasma concentration data was processed by DAS 2.0 software. The main pharmacokinetic parameters were statistically analyzed by SPSS 17.0 software. Results The encapsulation efficiency of the prepared micelles was (84.43^+/-^2.93) %, the drug loading was (2.63^+/-^0.91) %, and the particle size was (63.75^+/-^4.12) nm. The release behavior of GEN micelles was in line with the Weibull model. The 6-month leakage rate of GEN micelles was 2.45%, and the drug loading decreased by 0.18%. The main pharmacokinetic parameters AUC0-t after GEN micelles and GEN emulsion 40 mg·kg^-1^ were injected into the tail vein of rats. They were (99.46^+/-^ 4.77) mg · L^-1^ ·h and (57.51^+/-^1.37) mg·L^-1^ ·h, and t1/2 were (7.48^+/-^1.15)h and (4.95^+/-^ 1.15)h, respectively, and Cmax was (16.03^+/-^1.20) mg·L^-1^ and (16.73^+/-^1.10) mg·L^-1^, CL are (0.36^+/-^0.02) L·h^-1^ ·kg^-1^ and (0.67^+/-^0.02)L·h^-1^ ·kg^-1^.

## Introduction

Spontaneously formed core-shell nanocarriers [1]. It has received extensive attention in the past decade due to its outstanding anti-blood dilution ability, long-circulating characteristics, ability to greatly increase the solubility of poorly soluble drugs, and slow and controlled release of drugs, and strong ability to penetrate tumor sites. Genstein (GEN) is an isoflavone compound extracted from the legume plant Sophora japonica L. Numerous studies have confirmed that GEN has estrogen, anti-oxidation and inhibition of topoisomerase activity, inhibition of protein tyrosine kinase activity, induction of programmed cell death, inhibition of angiogenesis, etc. [2], currently a new class of drugs “dye wood” “Capsule” has entered Phase II clinical research and is a promising cancer chemopreventive and therapeutic agent. However, GEN is almost insoluble in water, and its direct oral bioavailability limits its clinical application. In this experiment, GEN copolymer micelles were prepared by MePEG-PLGA with good biocompatibility and application safety to improve the bioavailability of the drug, increase its targeting and improve the therapeutic effect. At the same time, its in vitro release, initial stability and pharmacokinetic behavior in rats after intravenous injection were investigated, which provided a theoretical basis for the development of clinical drugs and new dosage forms. 1 Instruments and materials 1.1 Instruments TECNAIG2 transmission electron microscope (Philips, the Netherlands); Mastersizer 2000 laser particle size analyzer (Malvern, UK); LC-2010AHT high performance liquid chromatography (Shimadzu, Japan); BT25S electronic balance (Sartorius, Germany); Scientz-IID ultrasonic cell crusher (Ningbo Xinzhi Biotechnology Co., Ltd.). 1.2 Material GEN reference substance (China National Institute for the Control of Pharmaceutical and Biological Products, content ≥ 98%, batch number: 111704-200501); GEN (Xi’an Xiaocao Plant Technology Co., Ltd., content: 98.19%, batch number: XC100310); MePEG-PLGA [Jinan Biotech Co., Ltd., Mr(MePEG)=4 000, Mr(MePEG): Mr(PLGA)=1:3, Mr(LA):Mr(GA)=75:25]; Dialysis Bag (Sigma, USA) Company, MW: 8 000∼14 400); Poloxamer 188 (BASF, Germany); Genistein intravenous emulsion (home-made, batch number: 20110712); methanol is chromatographically pure, water is deionized water, and the rest of the reagents are of analytical grade. 12 clean Wistar rats, each half, body weight (200^+/-^20)g, provided by GLP Laboratory, Heilongjiang University of Traditional Chinese Medicine, Laboratory Animal License No.: SCXK (Black) 2008-0004

## Results and Methods

The micelles were prepared by modified self-emulsifying solvent diffusion method (modifiedSESD) [3], and 21 mg GEN, 110 mg MePEG-PLGA were weighed into an Erlenmeyer flask, and acetone:ethanol = 3:2 mixed organic solvent 10 mL was added. The water bath was completely dissolved at 50 °C. The resulting mixed solution was dropped into a pre-formulated 20 mL aqueous solution containing 1% Poloxamer 188 at a rate of 3 drops of ^-^s ^-1^ under mechanical agitation at 500 r^-^min^-1^. After the completion of the titration, stirring was continued for 10 min, and the obtained product was removed under reduced pressure at 40 ° C to remove the organic solvent, filtered through a 0.45 μm microporous membrane, mixed with 3% trehalose, and stored after lyophilization.

Take GEN micelles lyophilized product, add water and shake gently to disperse and then add dropwise on copper network, negative staining with 2% sodium phosphotungstate solution, observe the particle morphology under transmission electron microscope, see Figure 1. It can be seen from the figure that after the micelles are formed into lyophilized injections, they are still spherical entities, and the shape is relatively round and the distribution is relatively uniform.

**Figure 1.**
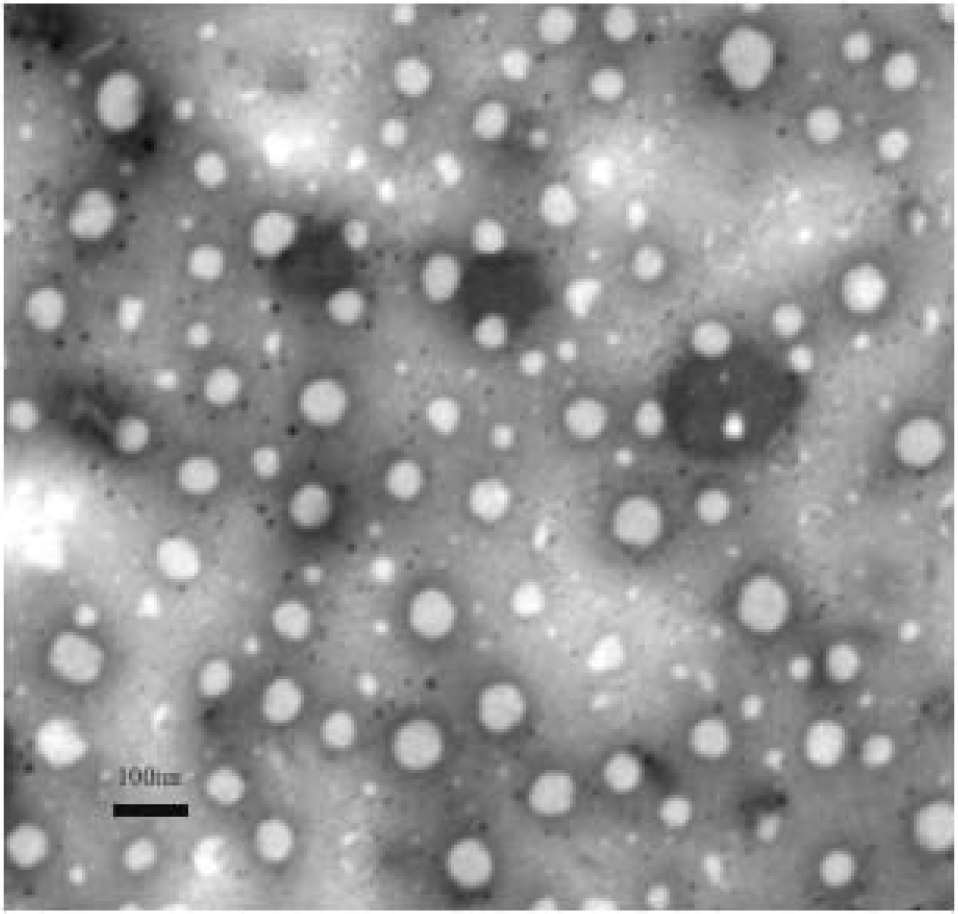
TEM images of micelles

**Figure 2.**
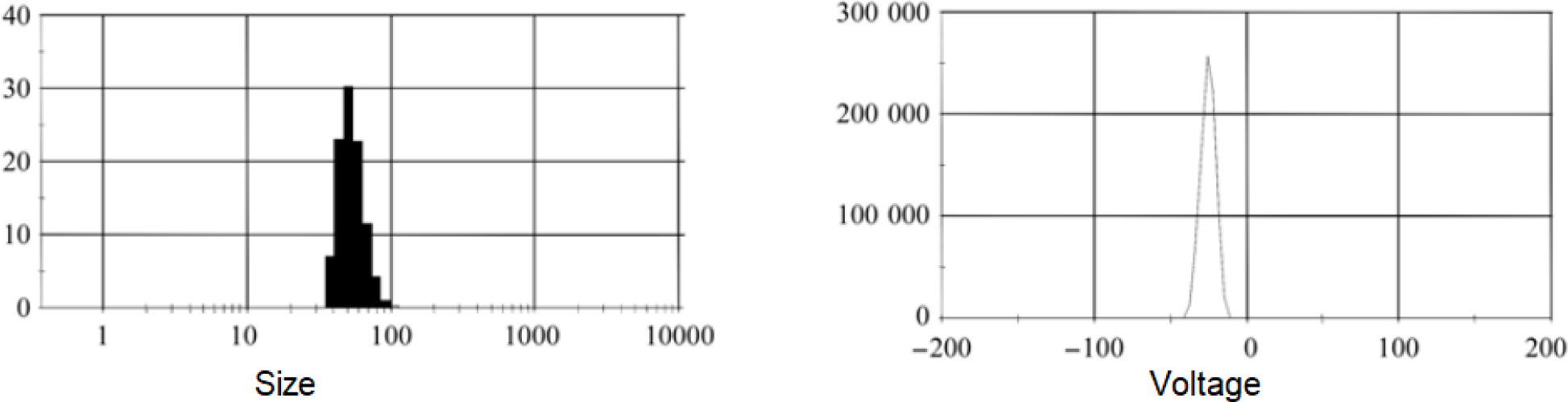
Micelle size and charge analysis

Encapsulation efficiency and drug loading of GEN micelles The GEN content was determined by reversed-phase HPLC. Chromatographic conditions: Diamonsil C18 column (4.6 mm × 250 mm, 5 μm), methanol-water (60..40), flow rate 1 mL·min^-1^, detection wavelength 260 nm, injection volume 10 □L. Under this chromatographic condition, the concentration of the GEN peak (Y) is linearly regressed to the concentration (X). The resulting standard curve equation is: Y=124 316X–38 008 (r=0.999 9), intra-day and inter-day precision RSD98 % indicates that the method meets the analytical requirements. Load 4 mL of GEN micelle solution into the treated dialysis bag, tie the bag tightly, and suspend it in a small cup containing 86 mL of release medium. Place the cup in a constant temperature water bath shaker at 37 °C. The speed (100 r·min^-1^) was shaken. At 360 min, 2 mL of the dialysate solution was accurately aspirated. After treatment, the amount of free GEN in the micelles was determined according to the above chromatographic conditions. At the same time, accurately extract the same volume of GEN micelle solution, accurately add 8 mL of acetonitrile, vortex for 1 min, filter with 0.45 μm microporous membrane and perform HPLC to determine the total amount of GEN contained in the micelle (W total). Calculate drug loading and encapsulation efficiency: ER% = (W total ^-^ W free) / W total × 100%; DL /% = (W total ^-^ W free) / W micelle 100% (where W is always The total amount of GEN contained in the GEN micelles, W is the amount of free GEN in the dialysate, and the W micelle is the sum of the amount of GEN and excipient contained in the micelle. The results are shown in Table 1.

Accurately add 4 mL of water for injection, disperse it, transfer it to the treated dialysis bag, tie the bag tightly, suspend it in a small cup containing 86 mL of release transmitter, and place the cup in a constant temperature water bath shaker at 37 °C. The constant speed in the water bath was shaken at 100 r^-^min^-1^. After the start of the release, take 1 mL of the drug release medium at 0.25, 0.5, 1, 2, 4, 6, 8, 10, 12, 24, 36, 48 h (add the same amount of isothermal blank medium at the same time). After filtration through a 0.45 μm microporous membrane, the filtrate was taken for injection and the cumulative release rate Q (%) of the drug was calculated. Accurately draw a quantity of GEN solution (GEN to release the medium as solvent, the content is equivalent to GEN micelle lyophilized product), put it in the dialysis bag, and test it in the same way. The sampling time was 5, 10, 20, 30, 45, 60, 90, 120 min, and the cumulative release rate Q (%) was calculated, and the in vitro release curve was plotted. The results are shown in Fig. 3.

**Figure 3.**
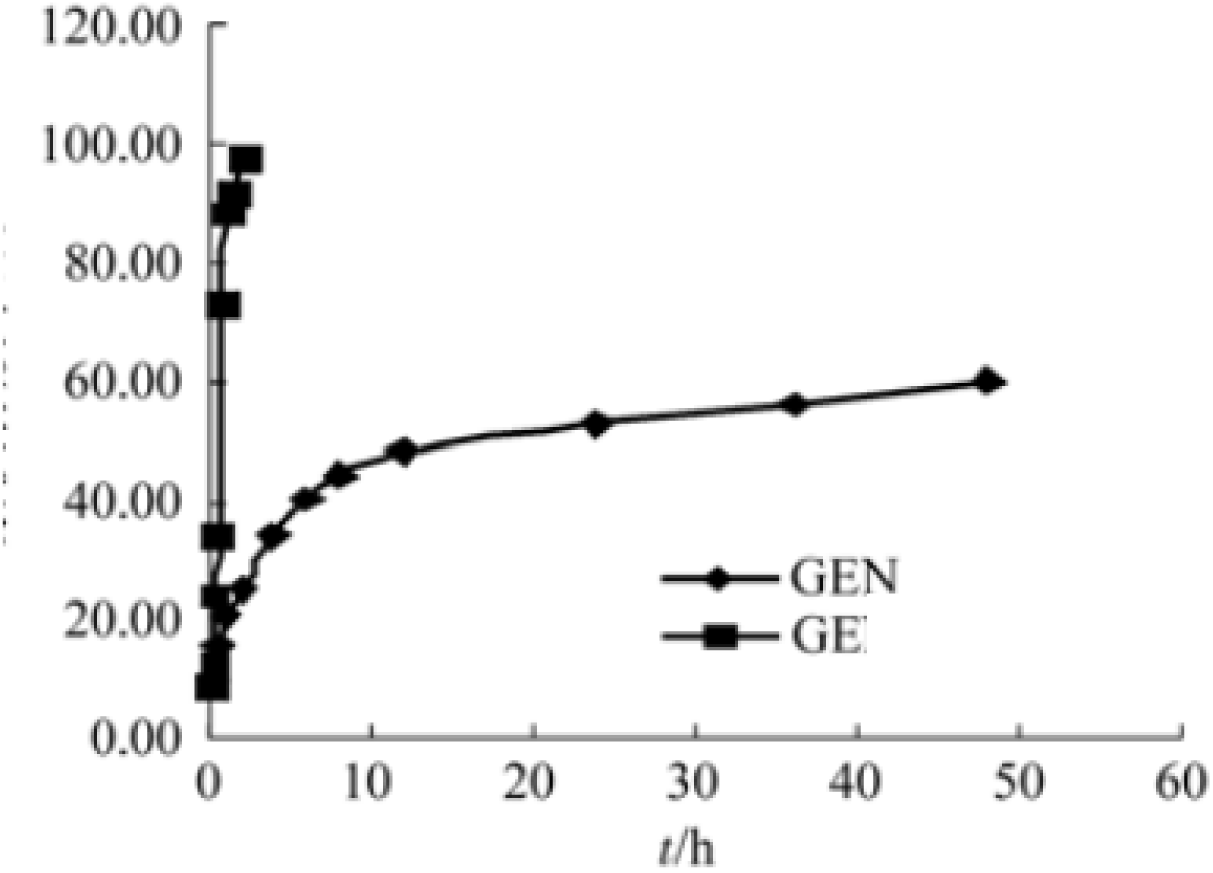
Releasing kinetic analysis

It can be seen from the results that the release of GEN drug substance is faster, and the basic release is complete at 2 h, while the release of GEN micelles is faster and the release is slower in the later stage, which has a significant sustained release effect compared with GEN drug substance. The first-order kinetic equation, Higuchi equation, Niebergull square root law, Hixcon-crowell cube root law, Ritgerpeppas equation and Weibull equation were used to fit the cumulative release percentage of GEN micelles at each time point, and the regression equation was obtained. The joint degree is judged by R2 and AIC values. The Weibull model has the best fitting effect. The equation is Ln[Ln(1/1-Q)]= 1.266Ln(t)4.768, R2=0.954, AIC=28.99. It has been confirmed that it has a certain sustained release property, which provides a deep theoretical basis for further in vivo research.

The prepared 3 batches of GEN micelles were stored in a refrigerator at 4 °C, and samples were taken after 1, 10, 30, 90, 180 days. After hydration, the encapsulation efficiency and drug loading were determined. Stability, the results are shown in Table 2. The results showed that the leakage rate of GEN micelles was 2.45% at 6 months and the drug loading decreased by 0.18%. The initial stability test showed good stability and achieved the expected purpose.

Accurately draw 100 μL of plasma sample into a 1.5 mL centrifuge tube, add 200 μL of methanol to precipitate plasma protein, vortex for 1 min, centrifuge at 10 000 r·min^-1^ for 10 min, and aspirate all supernatant. Pass through the microporous membrane (0.22 μm) and test according to the chromatographic conditions under “2.4”. 2.7.2 Specificity examination The chromatograms of the plasma samples collected after 1 hour of blank plasma and intravenous GEN micelles are shown in Figure 4. It can be seen from the results that the retention time of GEN is 11 min, the baseline noise is small, and the metabolite peaks and other endogenous substances in the plasma do not interfere with the determination of GEN, indicating that the method has high specificity.

**Figure 4.**
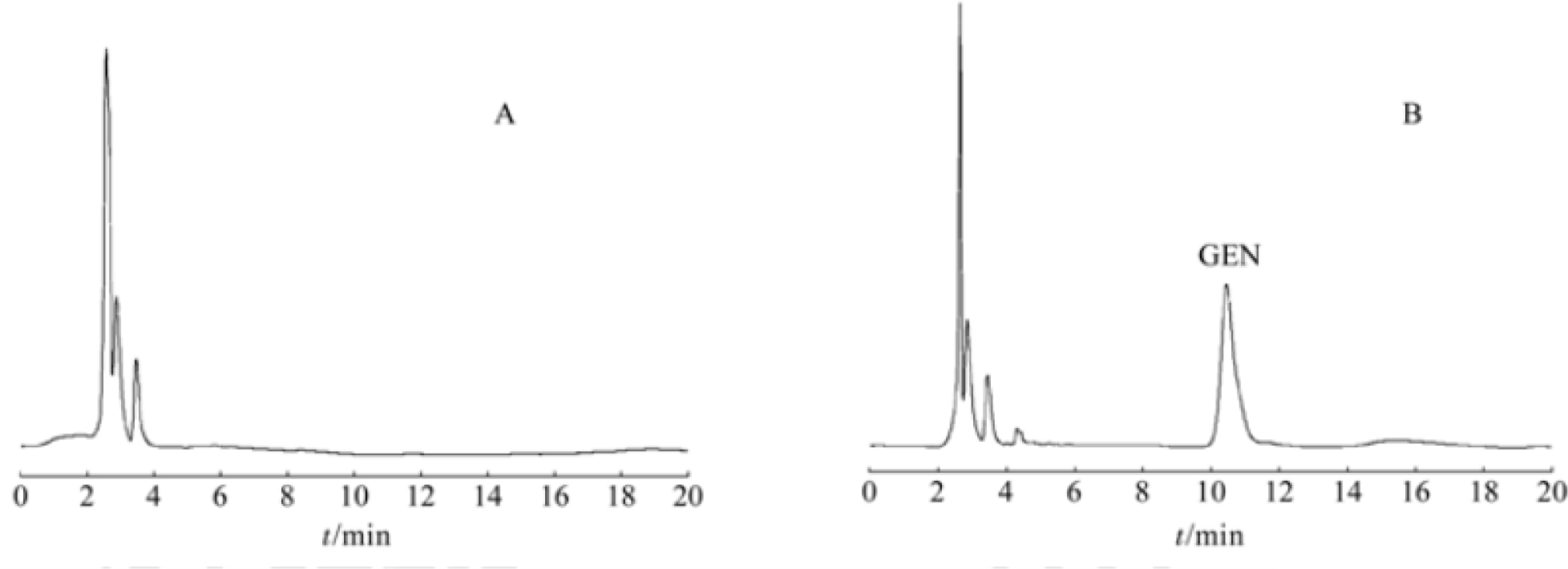
Serum retention time analyzed by HPLC.

Standard curve and linear range accurately absorb 200 μL of rat blank plasma, and add 20 μL of different concentrations of GEN standard series solution to prepare GEN blood concentration of 0.1, 0.4, 2, 8, 24, 32, 40 μg The plasma sample of mL^-1^ was treated according to the method of “2.7.1” and the sample was injected according to the chromatographic conditions under “2.4”. Taking the concentration of GEN in the plasma as the abscissa and the peak area as the ordinate, linear regression was performed to obtain a linear regression equation: Y=90 749X+1 779, r=0.999 9. The results showed that GEN was linearly good in 0.1∼40 μg·mL^-1^. The minimum detection limit is 0.02 μg·mL^-1^ in terms of S/N=3.

Take three low-, medium-, and high-quality (0.1, 8, 40 μg·mL^-1^) quality control samples, and continuously inject five samples per sample to calculate the intra-day precision; the same for 5 consecutive days. Conditional injection, calculation of daytime precision. The results show that the method’s daytime and intraday precision RSD are both

Weigh accurately 0.60 g of soybean lecithin and 5 g of soybean oil for injection in a beaker, stir at a constant temperature of about 10 °C for about 10 min, add GEN 500 mg, stir with stirring to reach the temperature. 70 °C and kept constant. Accurately weigh 1.25 g of medical glycerin and Poloxamer 1 881.00 g in a beaker, add 45 mL of water for injection, stir with stirring, and continue heating and stirring to keep the temperature at 70 °C and keep it constant. The oil phase was slowly added to the aqueous phase at a constant temperature of 70 ° C and continuously stirred, and stirring was continued for 10 min to obtain colostrum. The colostrum was prepared by ultrasonic ultrasonic cell pulverizer ice bath probe for 15 min (ultrasonic 2 s, intermittent 2 s), power 750 W, and GEN intravenous emulsion was prepared. The GEN intravenous emulsion prepared by the above method was milky white, uniform in appearance, no wall hanging phenomenon, no particles and precipitates were precipitated, and stored in a refrigerator at 4 ° C for two months without delamination and flocculation, and the state was good. It was observed by transmission electron microscopy (TEM) as a spherical shape with uniform particle size. The average particle size was 232 nm and the Zeta potential was −21.1 mV with a laser scattering particle size distribution and a Zeta potential meter. The drug loading was 1 mg· The mL^-1^, centrifugal acceleration test (4 000 r·min^-1^) showed good stability^1-14^.

Experimental design Rats were randomly divided into 2 groups, 6 in each group, and each half. Fasting for 12 h before the experiment, free to drink water. According to the literature [5-6] and pre-test, the doses were determined to be GEN emulsion group (40 mg·kg ^-1^), GEN micelle group (40 mg·kg ^-1^, and two bottles of GEN micelle lyophilized product)., equivalent to GEN 4.0 mg, disperse with 1 mL of water for injection before use. After weighing each group, the tail vein is administered intravenously, 0, 0.083, 0.17, 0.5, 1, 2, 4, 6, after administration. At 8, 10, 12, 24 h, about 0.4 mL of blood was taken from the posterior venous plexus. The blood sample was placed in a 1.5 mL heparinized centrifuge tube, centrifuged at 10 000 r^-^min^-1^ for 10 min, and the average drug-time curve of the supernatant of the 100 μ GEN emulsion group and the micelle group was shown in Fig. 5. From the results, the blood concentration-time curve of the GEN micelle group was more stable than that of the GEN emulsion group. Except for the initial blood concentration value, the blood concentration of the GEN micelle group was significant at each time point. Higher than the emulsion group, and still have a higher concentration distribution at 24 h. Since the cells at the tumor site have sufficient blood supply, the permeability is hyperactive, and the higher the concentration of the drug in the blood, the longer the maintenance time, the more opportunities for the drug and the tumor tissue to contact, and the more favorable the drug is delivered to the tumor tissue. Therefore, GEN micelles help to improve the anti-cancer effect of GEN.

**Figure 5.**
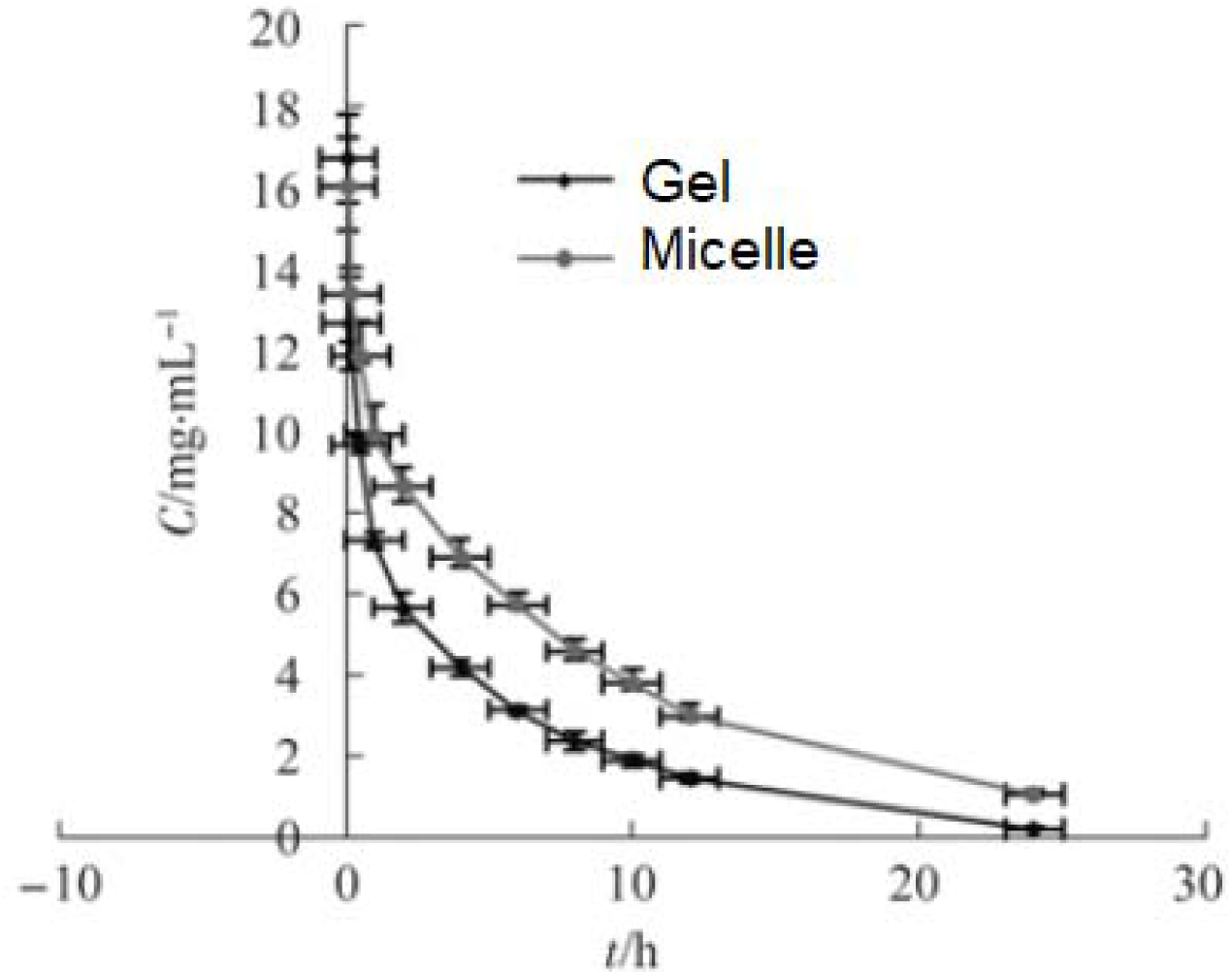
Pharmacokinetics comparison between gel and micelle.

The DAS2.1 software was used to fit the non-compartment model and calculate the pharmacokinetic parameters. The peak concentration (Cmax) and the peak time (Tmax) were measured and the area under the curve (AUC0^-^t) was calculated by the trapezoidal method. The main pharmacokinetic parameters (AUC0^-^t, AUC0^-^∞, t1/2, CL, MRT0^-^t) of the GEN emulsion group and the micelle group were tested by SPSS 17.0 software. The results are shown in Table 3. The results showed that there were significant differences in AUC0^-^t, AUC0^-^∞, t1/2, and CL between the GEN emulsion group and the micelle group at the same dose (P<0.05).

## Discussion

GEN is almost insoluble in water, which limits its application and research. Therefore, we must first try to improve the solubility of the drug and improve its bioavailability. The classical method of solubilization in pharmaceutics mainly uses a mixed solvent or a solubilizing agent, etc., but these are either unsatisfactory or have a large toxicity. For example, polyoxyethylene castor oil EL which solubilizes cyclosporine A and paclitaxel can cause side effects such as allergies, hypertension and neurotoxicity [7]. The commonly used surfactant Tween-80 can cause allergic reactions such as allergies and hemolysis [8], cholate is irritating to the mucosa, causing pain and hemolysis when injected. As a drug carrier with great development potential, polymer micelles have basically overcome the above shortcomings. The phase II clinical study of paclitaxel micelle preparation Genexol-PM showed that only 4 of 69 patients developed allergic reactions [9]. The modified self-emulsifying solvent diffusion method (modified-SESD) was first reported by Murakami et al. for the preparation of nanoparticles. The organic phase used a mixed solution of acetone and ethanol instead of the mixed dichloromethane and acetone commonly used in the SESD method. The solution, wherein the ratio of acetone to ethanol in the organic phase has a significant effect on the size of the emulsion droplets, this method overcomes the disadvantages of using toxic organic solvents (such as dichloromethane, chloroform, etc.) in the preparation process, and does not require high-energy equipment (such as Quality, ultrasound, etc.), providing the possibility of mass production. However, since the polyvinyl alcohol (PVA) used in the original method may be carcinogenic and difficult to remove from the surface of the nanoparticles, this method has been further improved by the subject, and replaced by the highly safe Poloxamer 188. PVA, the prepared micelle has a smooth surface, a round shape, and a high encapsulation efficiency and a small particle size.

The encapsulation efficiency of the micelle prepared by this subject is more than 80%, but the drug loading is relatively low. The reason may be that the smaller particle size limits the amount of drug contained in the hydrophobic core, so it is necessary to further study it. GEN micelles for higher encapsulation and drug loading. It has been reported in the literature [10-11] that this problem can be solved by covalently bonding the drug to PLGA. Therefore, this topic will further attempt to bond the drug to the carboxyl group at the end of PLGA, and then prepare GEN micelles.

Blood samples contain large amounts of protein, and direct injection can block the column and increase the column pressure, so proper pretreatment is required to remove the protein. Currently commonly used methods include protein precipitation (PPT), liquid-liquid extraction (LLE), solid-phase extraction (SPE), and the like. The author first tried to use methanol and acetonitrile as a precipitating reagent PPT method, and found that using methanol as a precipitant can get better results, and the PPT method is simple and easy, so methanol is selected as a precipitant to remove protein.

Since there is no GEN injection preparation available at home and abroad, and GEN is almost insoluble in water, it cannot be directly dissolved in physiological saline for injection into the tail vein of rats. According to the principle of reference preparation selection, GEN is used as an emulsion in the experiment. Compare the formulation with GEN micelle lyophilized injection. Compared with GEN emulsion, the elimination half-life (t1/2) and average residence time (MRT0^-^t) of the micelle group were significantly prolonged, mainly because the hydrophilic PEG was covalently attached to the PLGA via the ester bond. The formation of PEG-coated nanoparticles can effectively avoid the recognition and phagocytosis of the reticuloendothelial system (RES), prolong the circulation time in vivo, and achieve long-acting sustained-release effects. In addition, the AUC of intravenous GEN micelles was greater than that of the emulsion group, which was also the result of a decrease in plasma clearance.

## Conclusion

The prepared GEN micelles have regular morphology, narrow particle size distribution, high encapsulation efficiency, certain sustained release characteristics, good stability, and obviously change the pharmacokinetic behavior of GEN, so that the elimination is slowed down. Increased bioavailability of the drug. Future efforts will focus on translating this promising research to the clinical settings.

